# Time-Restricted Feeding Attenuates Kidney Damage and Preserves Renal Function in Mouse Model of Chronic Kidney Disease

**DOI:** 10.1101/2025.09.08.674986

**Authors:** Terry Lin, Carlos Rey-Serra, Jessica Tituaña, Pablo Cannata, Hiep Le, Aidan Glina, Debashis Sahoo, Santiago Lamasp, Amandine Chaix, Satchidananda Panda

## Abstract

Chronic kidney disease (CKD) is a highly disabling and potentially deadly condition for which there is no cure. With renal damage risk factors such as hypertension, metabolic syndrome and type 2 diabetes on the rise, the prevalence of CKD is increasing worldwide. New therapeutic approaches to CKD management are thus warranted. Time-restricted feeding, a dietary intervention in which daily food intake is limited to a consistent time window, has shown benefits in the context of metabolic disease management. Boolean implication network model of human CKD gene expression data and data from mouse TRF kidney implied TRF could attenuate kidney injury from CKD. We tested the effect of TRF in mouse models of kidney damage under high-fat high-sucrose feeding to induce a metabolic-disease prone environment. Using gold standard pre-clinical models of renal fibrosis, we discovered that TRF protected from kidney damage and clinical features of CKD. At the molecular level, the effects of TRF were pleiotropic with benefits in pathways involved in renal inflammation, fibrosis, and ER stress. Importantly, our results also suggest that TRF can confer early protection from metabolic alterations implicated in kidney damage.

## Introduction

Chronic kidney disease (CKD) affects more than 10% of adults worldwide and its prevalence is growing. In 2016, CKD was the 16^th^ leading cause of death and is projected to be the 5^th^ by 2040 (Francis et al., 2024). This represents a worldwide clinical challenge as CKD is a highly disabling disease, with a high mortality rate (4.6% of total mortality (Collaboration, 2020) and thus far irreversible damage. The underlying causes of CKD are multifaceted, yet diabetes and hypertension remain the predominant risk factors. It is estimated that 40% of diabetic patients develop CKD and that many cases of nephro-angio-sclerosis are related to hypertensive disease (Anders et al., 2018; Hall et al., 2014; Kazancioglu, 2013). Beyond strategies to maintain or restore proper blood pressure and glycemic controls, therapeutic options to prevent the progression of CKD are very limited. Furthermore, anti-hypertensive and diabetic treatments are insufficient to halt the progression of CKD towards renal failure, urging the development of new approaches to attenuate CKD progression.

Renal tubulointerstitial fibrosis is a convergent late-stage pathological outcome observed across many CKD etiologies. Renal fibrosis is characterized by the aberrant accumulation of extracellular matrix (ECM) proteins, partially attributed to unresolved chronic tissue inflammation (Djudjaj & Boor, 2019; Duffield, 2014; Humphreys, 2018). However, the metabolic rewiring in tubular epithelial cells is also a major mechanism underlying the development of renal fibrosis. Recent studies have demonstrated a key role of diminished fatty acid oxidation (FAO) during renal fibrosis and hence the potential to combat kidney fibrosis by restoring this metabolic pathway (Kang et al., 2015; Miguel et al., 2021).

The circadian clock also plays a critical role in the regulation of metabolic homeostasis via the central control of nutrient uptake and behavior through the cell-autonomous, diurnal coordination of metabolism-related gene expression (Panda, 2016; Reinke & Asher, 2019). Dampened circadian rhythms have often been associated with metabolic dysfunction and increased risk of metabolic disease (Longo & Panda, 2016). This highlights the importance of a robust circadian system in maintaining metabolic health. In the context of chronic metabolic disease associated with obesity, nutritional interventions can be efficacious in reinforcing circadian rhythms by delaying and/or preventing the development of these comorbidities. Importantly, enforcing feeding-fasting rhythms by time-restricted feeding (TRF) was shown to prevent and even reverse obesity-related metabolic syndrome (Chaix et al., 2014; Hatori et al., 2012). Specifically, in both animal studies and in pilot human studies, TRF can prevent or reduce the severity of glucose intolerance, hypertension, and fatty liver diseases (Manoogian et al., 2022) – all three of which are also known to increase the risk for CKD. Particularly in the liver, TRF was shown to enhance FAO, reduce fibrosis, and reduce ER stress secondary to an obesogenic diet. TRF also reduced systemic and tissue-specific inflammation, both of which are key factors for fibrosis development. (Chaix et al., 2019; Chaix et al., 2014; Hatori et al., 2012). Hence, there is a sound premise that TRF would be effective in the context of kidney injury, limiting inflammation and halting the progression of CKD.

Mouse models of chronic kidney disease are often accelerated when compared to human disease, but the pathogenesis and consequences of kidney damage are well paralleled. Using data from previous TRF studies, we hypothesized that similar metabolic and anti-inflammatory effects in the kidney under TRF regimen would limit renal fibrosis development and protect from maladaptive kidney dysfunction. Using gold standard pre-clinical models of renal fibrosis, we discovered that TRF protects from kidney damage and clinical features of CKD following both chemical and surgical insults. The effects of TRF were pleiotropic with benefits in pathways involved in renal inflammation, fibrosis, and ER stress. Importantly, our results also suggest that TRF can confer early protection from metabolic alterations implicated in kidney damage while retaining kidney function.

## Methods

### Time- restricted feeding protocol

All animal experiments were carried out in accordance with protocols reviewed and approved by the IACUC of Salk Institute. Eight-week-old male C57BL/6J mice fed *ad libitum* with normal chow were previously adapted to a 12h light: 12h dark cycle for 2 weeks before randomly assigned to *ad libitum* (AL) or time-restricted feeding (TRF) group and fed a 45 kcal% high fat diet (D12451, Research Diets, New Brunswick, NJ, USA). TRF mice had access to food for 9 h during the dark phase, from ZT13 to ZT22, where ZT0 indicated light on and ZT12 light off. Food intake and body weight were monitored every week, and body composition was analyzed after 6 weeks using a body composition analyzer (EchoMRI^TM^-100H). At the end of each experiment, animals were euthanized at approximately ZT13-16.

### Mouse models of chronic kidney disease

#### Unilateral ureteral obstruction (UUO)

UUO procedure was performed as described previously (Chevalier et al., 2009; Rey-Serra et al., 2023). The surgical procedure was conducted on a sterile bench using aseptic techniques. Mice were anesthetized with 2% isoflurane (depth confirmed by toe pinch), and eye lubricant was applied to eyes to prevent corneal drying. 0.5 mg/kg of buprenorphine was given subcutaneously for analgesic purposes. The hair in the abdominal area was shaved by wetting the hair with a combination of warm water and soap and then shaving with a scalpel. The underlying skin was cleaned with betadine followed by ethanol. An incision was made in the abdominal wall. Then, the left kidney was isolated to reveal the ureter and two ligatures were made, one near the kidney and one near the bladder. The intermediate ureter was removed, and the kidney was returned gently to its place. Finally, the abdominal incision was closed with sutures. Mice were euthanized by CO_2_ overdose 3 days after UUO. Immediately after the sacrifice, blood samples were collected by cardiac puncture and kidneys were collected after perfusion with phosphate buffered saline. Contralateral kidneys were used as controls.

#### Folic acid nephropathy (FAN)

FAN was performed as previously described (Fink et al., 1987; Rey-Serra et al., 2023). Mice were given 250 mg/kg of folic acid intraperitoneally (i.p) (Sigma-Aldrich) dissolved in 0.3 M sodium bicarbonate (vehicle) and control mice were given 0.1 ml of the vehicle. Mice were sacrificed 15 days after folic acid injection. Blood and kidney samples were collected as indicated in the UUO procedure.

### Assessment of kidney function

Blood urea nitrogen (BUN) and creatinine were analyzed in serum samples by using the BUN Colorimetric Detection Kit and the QuantiChrom^TM^ Creatinine Assay Kit (Ann Arbor, MI), according to manufacturer’s instructions.

### Histological and immunohistochemical analyses

Kidney samples were fixed in 4% neutral buffered formalin before being embedded in paraffin and cut into 3 or 5 µm tissue sections. Hematoxylin and Eosin (H&E) and Sirius Red stains were performed on 5 *µm* sections using standard procedures. 3 µm sections were deparaffinized for IHC and antigen retrieval was performed with 10 mM citrate sodium buffer by using the PT Link (Dako, Santa Clara, CA, USA). Endogenous peroxidase and non-specific protein binding sites were blocked with 3% H_2_O_2_ and with 4% bovine serum albumin (BSA) in 1X EnVision wash buffer (Dako, Santa Clara, CA, USA), respectively. Incubation with the primary antibodies TFAM (PA5-68789, ThermoFisher Scientific, Massachusetts, USA), KIM1 (AF1817, R&D systems, Minnesota, USA), αSMA (α-Smooth Muscle Actin (D4K9N) XP® Rabbit mAb #19245, Cell Signaling) and F4/80 (F4/80 (D2S9R) XP® Rabbit mAb #70076, Cell Signaling, Massachusetts, USA) was carried out overnight at 4°C. Incubation with biotinylated goat anti-mouse or anti-rabbit IgG secondary antibodies were performed for 1h at 4°C. VECTASTAIN ABC *Kit* (Vector Laboratories, Burlingame, CA, USA) was used for detection of biotinylated-secondary antibodies. Tissue sections were treated with 20 *µl/ml* 3,3’-diaminobenzidine (DAB, Dako) and counterstained with hematoxylin. Slides were scanned using the Ultra-Fast Scanner (Phillips, The Netherlands) and images were taken at 40x using the NDP.view2 software (Hamamatsu Photonics K.K, Japan). The intensity of Sirius Red (collagen deposition) and IHC was quantified automatically with the Image-pro Plus software (Media Cybernetics). Data was represented as a percentage of the ratio of positive area to total tissue.

### Histological evaluation of renal biopsies

H&E and PAS kidney sections were blinded and were assessed by a pathologist by using the Hill score (Hill et al., 2000). The following morphologic variables were evaluated: interstitial inflammation (Intinf), epithelial cells in the tubular lumen (EpLum), macrophages in the tubular lumen (MacrLum), tubular flattening (TubFlat), tubular necrosis (TubNec), tubular activity (TubAct), tubular pyknosis (TubPyk), tubular atrophy (TA) and interstitial fibrosis (IF). The lesions were classified on a scale of 0 to 3; where 0 referred to the absence of lesion, 1 to the lesions involving up to the 25% of the area analyzed, 2 to lesions involving 25 to 50% and 3 to lesions involving more than 50% of the area.

### RNA isolation and qPCR

The RNA from kidneys was isolated using a phenol-chloroform RNA extraction protocol. qScript® cDNA SuperMix (Cat No. 95048) was used for the reverse transcription of 1μg of total RNA, following manufacturer’s instructions. qRT-PCR was performed in triplicates using the iQBR^TM^ Green Power up (ThermoFisher Scientific, Massachusetts, USA) and the primers described in Supplementary Table 3 in QuantStudio5 (ThermoFisher Scientific, Massachusetts, USA). Relative mRNA expression was calculated using the ΔΔCt method and the mRNA levels were normalized to 18S.

### RNA sequencing and analysis

cDNA libraries were prepared using 500 ng of kidney RNA from 7-week TRF vehicle and FAN-injected mice with TruSeq Stranded mRNA kit (Illumina) as per manufacturer’s instructions. The libraries were quantified using Quant-iT™ dsDNA HS Assay Kit (ThermoFisher Scientific), pooled, and sequenced on the NovaSeq platform with PE100 coverage at the IGM Genomics Center, University of California, San Diego, La Jolla, CA. Libraries were individually mapped to the mm10 UCSC genome annotation and quantified using STAR (v2.5.3a) (Dobin et al., 2013). Data normalization and differential gene expression analysis was carried out via edgeR (v3.26.7)(Robinson et al., 2010). Over-representation analysis was performed using R package clusterProfiler (v4.7.1.003) and Metascape using ranked gene list, p-value < 0.05, in both KEGG and GO databases (Yu et al., 2012; Zhou et al., 2019). Gene set enrichment analysis (GSEA v4.2.3) was performed between FAN AL and FAN TRF using C2 KEGG, reactome, and GO biological processes databases from MSigDB, number of permutations set to 1000, with gene sets between 30 and 300 (Subramanian et al., 2005). This analysis was imported into Cytoscape (v3.9.1) using its EnrichmentMap (v3.3.3) and AutoAnnotate (1.3.5) applications(Reimand et al., 2019; Shannon et al., 2003). Venn diagrams were created using ggVennDiagram (v1.2.1), heatmaps were created using pheatmap (v1.0.12), dotplots were created using ggplot2 (v3.3.6).

Deconvolution of bulk RNA-seq of 7-weeks FAN plus TRF kidneys was done using R package MuSiC (v1.0.0) (Wang et al., 2019). Using publicly available single-cell RNA-seq references (GSE107585) (Park et al., 2018), MuSiC utilizes cell-type specific gene expression from single cell references to estimate cell proportions in bulk RNA-seq.

### Boolean analysis

Publicly available gene expression dataset GSE99340 (Shved et al., 2017) on human chronic kidney disease were downloaded from National Center for Biotechnology Information Gene Expression Omnibus website (NCBI-GEO). Boolean Analysis was conducted on a subset of 165 microdissected glomeruli samples obtained from human renal biopsies, which were profiled using the Affymetrix U133A microarray platform, selected from the dataset GSE99340. Mouse kidney TRF samples were taken from (GSE190389) consisting of 48 kidneys profiled from young male C57BL6 mice fed a 45% fat, Western diet (Deota et al., 2023). Boolean analysis was conducted as previously described (Sahoo et al., 2008). Briefly, expression values of each gene in this dataset were converted into a binary “low” or “high” expression value using a step function computed via the StepMiner algorithm (Sahoo et al., 2007). Boolean implication analysis determined statistically significant relationships between any two genes in this dataset based on their binarized expression patterns. Genes that showed equivalent expression patterns to each other were discovered and clustered into nodes via the Boolean Network Explorer framework (BoNE) (Sahoo et al., 2021). These nodes were connected together with the predominant non-equivalent Boolean relationships between the nodes (node A expression is low ⇒ node B expression is low, A low ⇒ B high, A high ⇒ B low, and A high ⇒ B high & A opposite to B) to create a Boolean implication network.

To compute the composite score, first the genes present in each cluster were normalized and averaged. Gene expression values were normalized according to a modified Z-score approach centered around StepMiner threshold (formula = (expr – (SThr+0.5))/3∗stddev). Weighted linear combination of the averages from the clusters of a Boolean path was used to create a score for each sample.

### Mitochondrial copy number determination

DNeasy Blood & Tissue Kit (Qiagen Cat. 69504) was used for the extraction of genomic DNA from kidneys according to the manufacturer’s instructions. Mitochondrial DNA copy number was determined using qPCR with unique mtDNA primers as described previously (Malik et al., 2016). Relative mtDNA copy number was represented as the mtDNA to nuclear DNA ratio.

### Statistical analysis

Data analyses were performed on GraphPad Prism 8.0 (GraphPad Software, La Jolla, CA, USA). Data are represented as mean ± standard error of mean (SEM). Statistical differences were determined with the non-parametric Mann-Whitney test when two independent groups were analyzed, while two-way ANOVA and Tukey’s post-test were used for more than two independent groups and two independent variables. A P-value of 0.05 was considered as statistically significant (*P<0.05, **P<0.01, ***P<0.001).

### Data availability

The kidney gene expression dataset is submitted to GEO.

To review GEO accession GSE245933: Go to https://www.ncbi.nlm.nih.gov/geo/query/acc.cgi?acc=GSE245933

Enter token efsxougyprqztov into the box. This token allows anonymous, read-only access to GSE245933 and associated accessions while they are private.

## Results

### Boolean implication analysis establishes an opposing relationship between chronic kidney disease (CKD) and time-restricted feeding (TRF)

Obesity and type 2 diabetes are associated with a systemic pro-inflammatory environment that contributes to progressive deterioration in renal function and eventually in the establishment of renal fibrosis and failure (Decleves et al., 2011). To establish whether our data collected in mice could inform the translational relevance/potential of TRF to human CKD, we correlated our preclinical data (GSE190389, 12-week-old C57BL/6J mice after 6 weeks of 45% Western diet *ad libitum* (FA) or time-restricted (FT), (Deota et al., 2023) with published human data (Shved et al., 2017). The bulk transcriptome of the kidney cortex was analyzed for differentially expressed (DE) genes between mice fed ad lib or TRF (Figure 1A). In the first analysis, the cortex samples that were collected as a timeseries (every 2hrs for 24hrs, (n=24/group)) were used as biological replicates to identify TRF effects independent of time. The top 25 DE genes are shown (Figure 1B). Given that there may be time-specific differences, we also performed a second DE analysis every 4hrs (n=4/group). The number of DE genes at these timepoints demonstrate a dramatic difference between the effect of TRF during the fed or fasted state of the mice (Figure 1C). We then performed GSEA enrichment of all DE genes from the timeseries analysis. Genes that were upregulated in TRF are involved in fatty acid metabolism, PPAR signaling, and mitochondrial transport while downregulated genes are involved in BCAA metabolism, angiogenesis, and lipid metabolism (Figure 1D-E).

**Figure 1.**
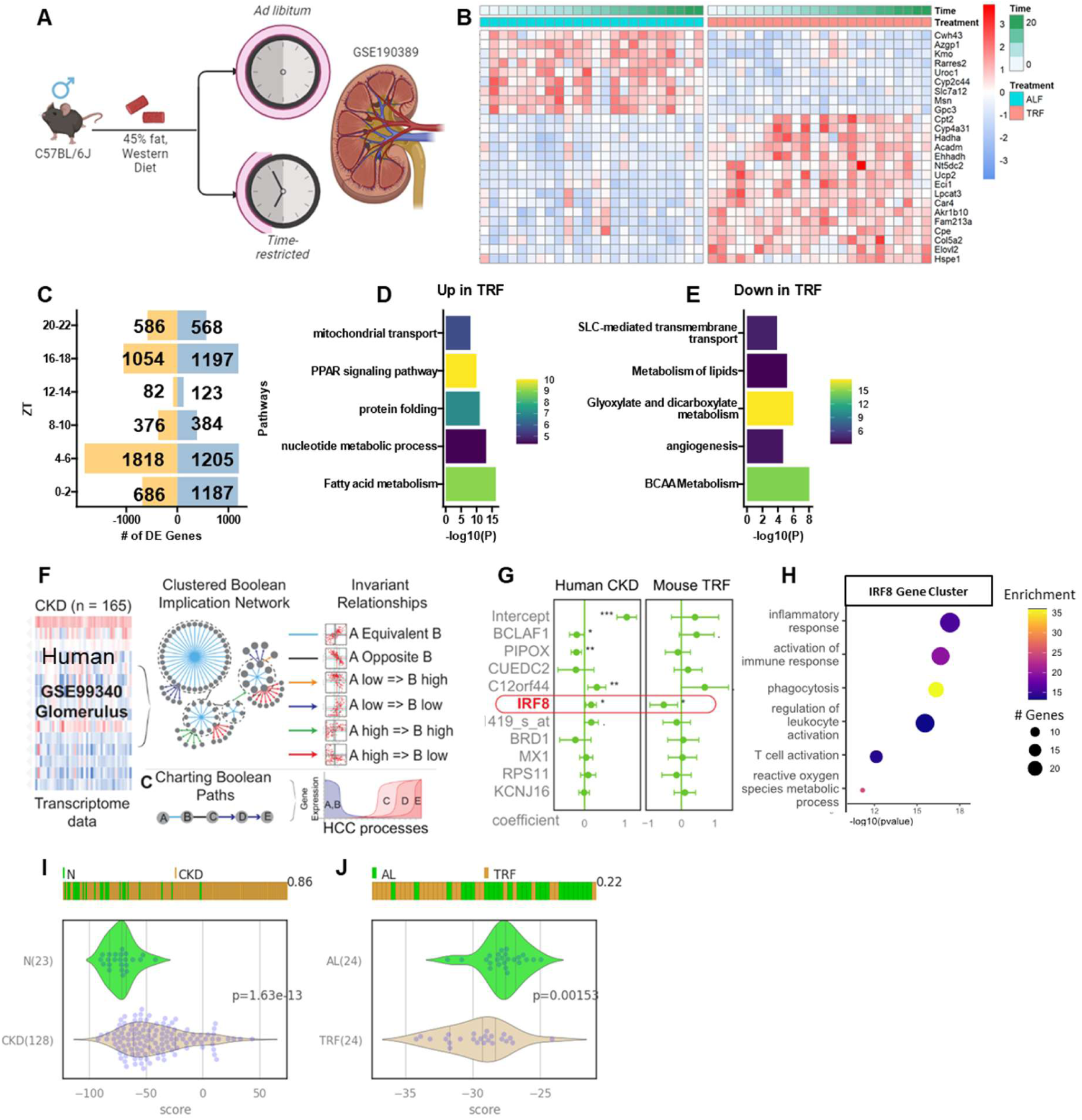
Effects of time-restricted feeding on kidney gene expression in mice. **(A)** Transcriptome obtained from kidneys of male mice on Western high-fat diet (GSE190389, n=48). **(B)** Heatmap of the top 25 genes differentially expressed between *ad lib* and time-restricted kidneys. **(C)** Number of DE genes up (blue) and down (gold) every 4 hours using 4 samples/group as biological replicates. Metascape pathway enrichment of DE genes upregulated **(D)** and downregulated **(E)** in TRF. **(F)** Visual representation of the construction of a Boolean implication network generated from human chronic kidney disease dataset (GSE99340, microdissected glomeruli samples, n=165). **(G)** Multivariate regression analysis of top 10 predictive equivalent clusters oriented by sizes for human CKD are tested for predicative ability in mouse TRF. **(H)** Top 6 GO pathways in human gene cluster IRF8. Violin plot of composite score of IRF8 gene cluster to predict sample identity in **(I)** human CKD (n=151) and **(J)** mouse TRF (n=48). In G, correlation coefficient with 95% confidence interval, *P<0.05, **P<0.01, ***P<0.001.

The mouse TRF kidney gene expression dataset indicated several pathways implicated in kidney disease were favorably modulated. Next, we asked for gene expression signatures of chronic kidney disease in humans and whether TRF has any specific effects on those expression signatures. First, we constructed a Boolean implication network using available chronic kidney disease datasets (GSE99340, microdissected glomeruli samples, n=165) from humans and TRF in mice (Figure 1F). Using an unbiased machine learning approach, Boolean analyses of human CKD samples found 5 significant gene cluster relationships with equivalent expression were found to be consistently altered (Figure 1G). Only one of these clusters (labeled as IRF8) is significantly downregulated by TRF treatment in mice. We selected this cluster of 69 equivalently expressed genes (labeled as cluster IRF8, Supplementary Table 1) that strongly correlated with human CKD gene signature and was downregulated by TRF treatment in mice (Figure 1G). Gene ontology analysis of the IRF8 cluster identified pathways positively related to immune/ inflammatory signatures (Figure 1H). Composite score of this gene cluster was able to reliably predict sample identity in human CKD (ROC-AUC: 0.86, p-value=1.63e-13) and mouse TRF (ROC-AUC: 0.78, p-value=1.89e-3) (Figure 1I-J). In summary, by utilizing publicly available datasets and machine learning, we identified potential gene-disease relationships targets that suggest TRF could prevent kidney inflammation and subsequent damage.

### TRF reduced kidney damage and preserved kidney function upon renal injury

To investigate the translational relevance of TRF in CKD, we tested the effects of TRF in mouse models of kidney injury in the context of a high-fat diet environment. Twelve-week-old male C57Bl/6J mice were fed a 45% high-fat 17% high-sucrose Western diet either *ad libitum* (AL) or TRF (9h food access in the dark phase ZT13-22) for 1 to 7 weeks to establish a diet-induced pro-inflammatory environment in the ad lib fed mice. As expected, and described previously (Chaix et al., 2014; Hatori et al., 2012), mice on TRF for 7 weeks gained less weight than mice on AL (12.9% TRF vs 26.7% AL) (Figure S1A), despite similar food consumption (Figure S1B). The TRF mice also showed a significant decrease in fat mass (2.53% TRF vs 11.2% AL) (Figure S1C).

Two different protocols of kidney injury were used: a chemical injury model induced by 250 mg/kg folic acid injection (Folic Acid Nephropathy/ FAN) and a surgery-based injury model by unilateral ureteral obstruction (UUO). In the first cohort, folic acid was injected 1 week (acute intervention model) or 7 weeks (sustained intervention model) after the start of the feeding protocol and renal outcomes were analyzed after 15 days (Figure 2A). In the second cohort, UUO was performed after 7 weeks of TRF, and outcomes measured 3 days post-surgery as an acute kidney injury model (Figure 2A).

**Figure 2.**
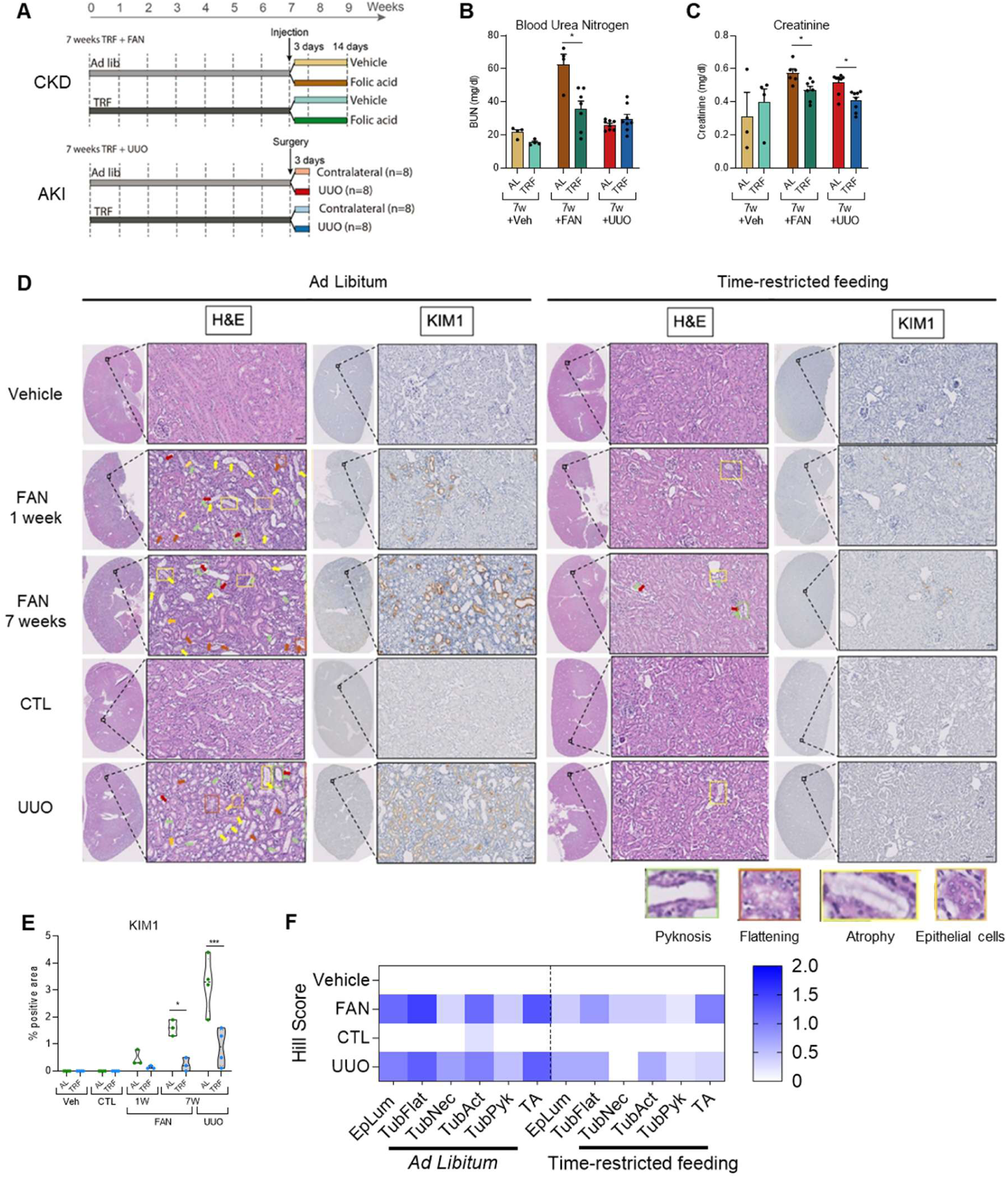
Time-restricted feeding prevents damage and kidney dysfunction in kidneys undergoing Folic Acid Nephropathy (FAN) and Unilateral Ureteral Obstruction (UUO). **(A)** Schematic of experimental design depicting TRF and models of nephropathy. **(B)** Blood urea nitrogen (BUN) and **(C)** plasma creatinine levels of vehicle, FAN, and UUO mice on 7-week TRF protocol. (n=4-8/group). **(D)** Representative microphotographs of H&E (left) and KIM1 IHC (right) staining of vehicle-treated and folic acid-treated kidneys from 1-week and 7-weeks TRF plus FAN, and contralateral/ obstructed kidneys from 7-weeks TRF plus UUO protocol (n=4-6/group). Arrows/rectangles denote: Tubular necrosis (red), Tubular pyknosis (green), tubular atrophy (yellow), epithelial cells in the tubular lumen (orange), tubular flattening (dark orange). Scale bar: 50 *µm*. **(E)** Quantification of KIM1 IHC in vehicle, contralateral, folic acid-treated, and UUO kidney sections, the data represent the mean ± SEM. *P<0.05, **P<0.01 compared with AL. **(F)** Heatmap of Hill score in AL and TRF kidneys blind-scored by pathologist. Morphological variables evaluated: epithelial cells in the tubular lumen (EpLum), tubular flattening (TubFlat), tubular necrosis (TubNec), tubular activity (TubAct), tubular pyknosis (TubPyk) and tubular atrophy (TA).

At the end of each study, we measured serum levels of blood urea nitrogen (BUN) and creatinine as proxy of kidney function. In the FAN model, both BUN and creatinine levels were significantly lower in mice on 7-weeks TRF than AL feeding (BUN: 35.8 mg/dL TRF vs 62.5 mg/dL AL, creatinine: 0.47 mg/dL TRF vs 0.57 mg/dL AL) suggesting improved kidney function in the TRF mice, while no differences were seen in the 1-week protocol (Figure 2B-C). Interestingly, we also observed significantly lower serum creatinine levels in mice under 7-weeks TRF subjected to UUO (0.41 mg/dL TRF vs 0.51 mg/dL AL) (Figure 2C). The lack of functional readouts is one of the principal disadvantages of the UUO model since the intact contralateral kidney is sufficient to preserve glomerular filtration rate and renal function. Thus, this result, together with the observed decrease in BUN levels in vehicle-treated mice after 7 weeks of TRF (15.4 mg/dL TRF vs 21.7 mg/dL AL) (Figure 2B), suggests enhancement of basal kidney function under TRF condition.

The extent of kidney injury/damage was then assessed using kidney histopathological scoring of H&E slides and analysis of KIM1 immunostaining. Both FAN and UUO models led to profound morphological alterations related to kidney injury (Figure 2D-F). Noteworthy, kidneys from TRF mice under both injury protocols showed less tubular atrophy (TA) and related morphological disturbances such as epithelial cells in the tubular lumen (EpLum), tubular flattening (TubFlat), tubular activity (TubAct), necrosis (TubNec) and pyknosis (TubPyk). Moreover, expression of the well-known renal damage marker KIM1 was significantly reduced in TRF kidneys irrespective of UUO or FAN kidney injury model (Figure 2D-E). Interestingly, TRF also reduced FAN-mediated injury in the acute 1-week protocol (Figure 2E) suggesting that early adaptations to TRF are sufficient to promote a protective environment against kidney injury. Taken together, these results show that TRF limits renal damage and dysfunction in the context of kidney injury.

### Kidney fibrosis and inflammation are reduced in mice under TRF

Since fibrosis is one of the main drivers of kidney dysfunction in CKD, we set out to determine the effect of TRF on kidney fibrosis. We analyzed kidneys from the FAN model since they were collected 2 weeks after the injection, a time point at which fibrosis is generally well-established in this model. Pathological examination of the histological slides revealed reduced interstitial fibrosis (IF) in folic-injected mice under TRF compared to ad libitum. This observation was confirmed with Sirius Red Collagen and Smooth Muscle Actin (αSMA) staining – two widely used markers of fibrosis. Quantification of the two stains show that both Sirius Red (4.2% TRF vs 13.6% AL) and αSMA (4.8% TRF vs 16.7% AL) (Figure 3A-C) are significantly lower in mice under TRF and observed both in the acute 1-week and prolonged 7-weeks long feeding protocol.

**Figure 3.**
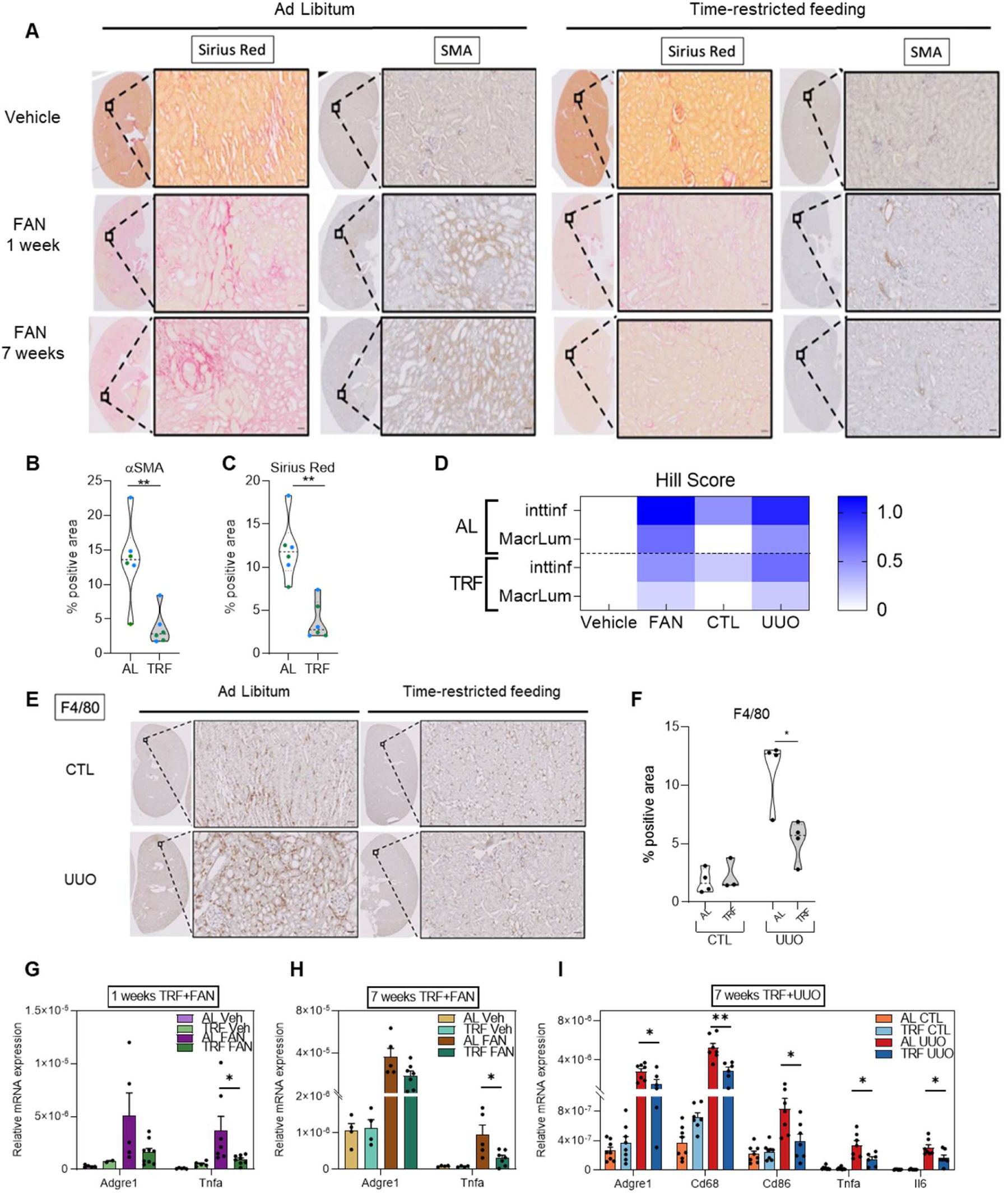
Time-restricted feeding reduces fibrosis and inflammation in folic acid- and UUO-induced kidney damage. **(A)** Representative microphotographs of Sirius Red (left) and α-Smooth Muscle Actin IHC (right) staining of vehicle and folic-acid treated kidneys from 1-week and 7-week TRF plus FAN, Scale bar: 50 *µm*. **(B)** Quantification of Sirius Red and **(C)** αSMA IHC in vehicle and folic acid-treated kidney sections from 1-week and 7-week TRF plus FAN (n=4-6/group). **(D)** Heatmap of Hill score in AL and TRF kidneys blind-scored by pathologist. Morphological variables evaluated: Interstitial inflammation (IntInf), Macrophages in the tubular lumen (MacrLum). **(E)** Representative microphotograph of F4/80 IHC immunostaining of contralateral and obstructed kidney in ad libitum and TRF mice from 7-weeks TRF plus UUO protocol, Scale bar: 50 *µm*. **(F)** Quantification of F4/80 IHC in kidney sections of 7-weeks TRF plus UUO protocol (n=4/group). mRNA expression of inflammatory genes in 1-week TRF plus FAN **(G)**, 7-weeks TRF plus FAN **(H)**, and 7-weeks TRF plus UUO **(I)**. The data represent the mean ± SEM. *P<0.05, **P<0.01, ***P<0.001 compared with their respective control kidney. #P<0.05, ##P<0.01, ###P<0.001 compared with AL.

We also assessed mRNA levels of fibrotic markers by qPCR in both the FAN and UUO model. The mRNA expression of the fibrotic markers *Fibronectin1, Col1a1, and Acta2* were significantly lower in TRF mice under all experimental conditions (1-week TRF + FAN, 7-weeks TRF + FAN, 7-weeks TRF + UUO (Figure S2A-C). Interestingly, the results observed in the UUO group just 3 days after surgery (Figure S2A) indicated protection by TRF even in early stages, when prevalent inflammation mediates the upregulation of pro-fibrotic genes. Furthermore, since ER-stress has been shown to favor fibrosis progression through the triggering of apoptosis (Kim et al., 2015) and was shown to be lowered in the liver under TRF (Wilson et al., 2020), we quantified mRNA levels of three key genes implicated in ER-stress response (*Chop, Xbp1, Bip*). There was a significant reduction in their expression in damaged kidneys from mice under 7-week of TRF, whereas no differences were seen in FAN kidneys from mice exposed to the short TRF protocol (Figure S2D-F).

Since inflammation also plays a critical role in the onset of tubulointerstitial fibrosis and TRF has been previously shown to reduce systemic and tissue-specific inflammation, we also analyzed the effect of TRF on kidney inflammation in both models of kidney injury. The histopathological analyses revealed a decrease in interstitial inflammation (IntInf) and macrophages in tubular lumen (MacrLum) in kidneys from TRF mice under both UUO and FAN models compared to AL (Figure 3D, S2G). Moreover, immunohistochemical evaluation of tissue inflammation 3 days after UUO using F4/80 antibody revealed a significant decrease in F4/80 positive stain (6.7% TRF vs 12.8% AL) in the obstructed kidney from mice under TRF compared to their AL counterparts (Figure 3E-F). Furthermore, mRNA levels of multiple macrophage markers (*Adgre1, Cd68, Cd86*) and pro-inflammatory cytokines (*Tnfa, Il6*) were also significantly decreased in kidneys from mice subjected to UUO after 7 weeks of TRF (Figure 3G-I). In all kidney injury models, significant differences were observed in F4/80 gene expression (*Adgre1*) analyzed by qPCR and mRNA expression of *Tnfa* was significantly lower in TRF mice compared to AL mice. TRF dampened the inflammatory and pro-fibrotic gene expression profile in response to injury and limited immune hyper-response and collagen deposition. Overall, these data support the role of TRF-mediated protective response against chronic inflammation and the establishment of kidney fibrosis.

### Transcriptomic analysis in kidney injury supports prevention of metabolic dysfunction by TRF

To understand the molecular changes accompanying the protective role of TRF on renal injury, we interrogated the global transcriptomic changes in kidneys from vehicle and folic acid-treated mice under TRF or AL for 7-weeks. For the data analysis, kidney samples were classified into 4 different groups to assess the specific effect of FAN, effect of TRF, and interactions: Vehicle-treated mice fed AL (Veh AL), folic acid-treated mice fed AL (FAN AL), vehicle-treated mice fed under TRF (Veh TRF) and folic acid-treated mice fed under TRF (FAN TRF). After data processing, a total of 17,836 genes were expressed in the kidney and considered for analysis. Differentially expressed genes with an adjusted p-value < 0.05 were considered statistically significant (Supplementary Table 2). Principal component analysis showed a high degree of gene expression changes upon folic-induced kidney injury with up to 66% of the variation in PC1 explained by treatment groups and a distinct separation between TRF and AL samples in injured kidneys (Figure 4A).

**Figure 4.**
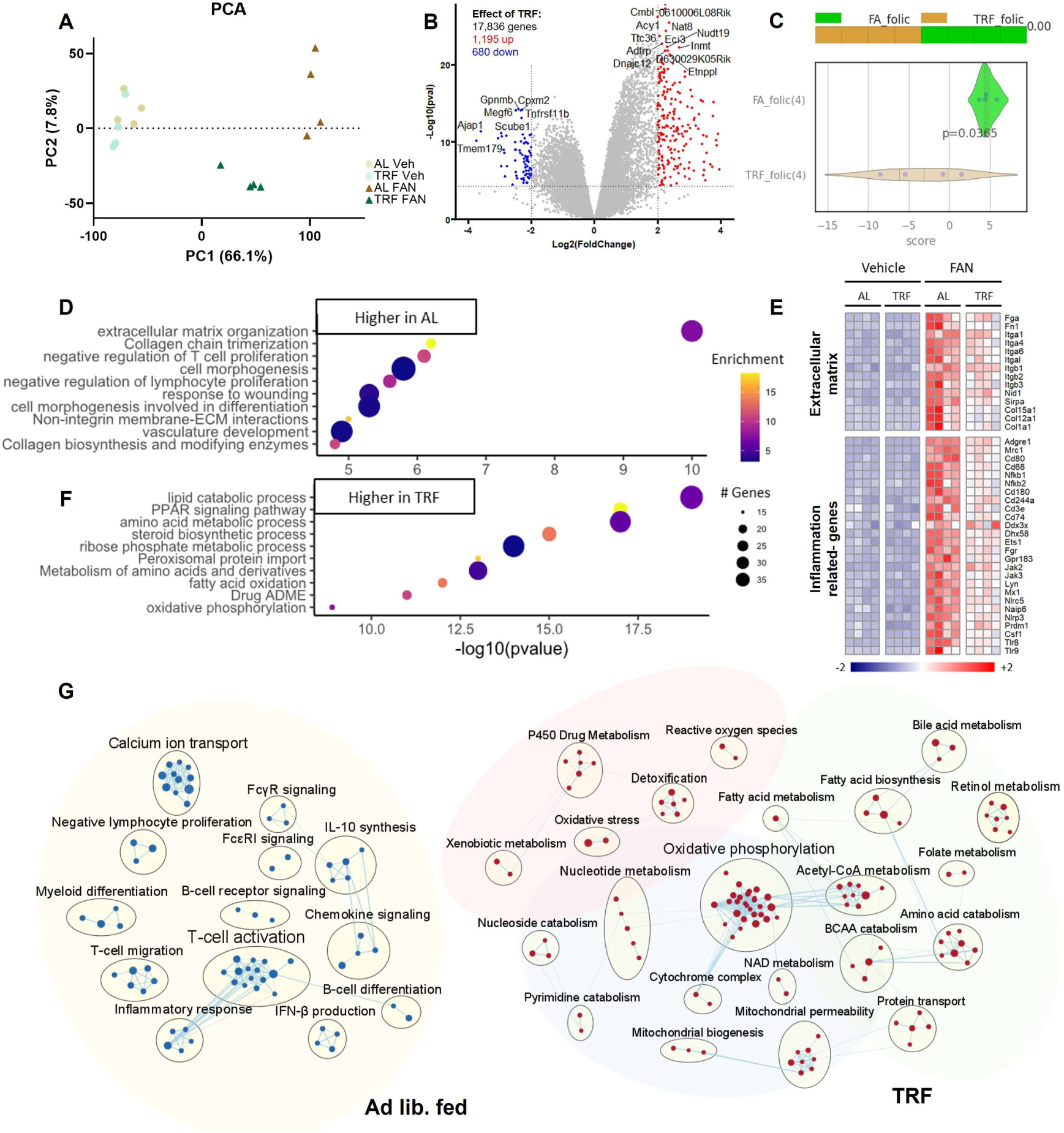
RNA-sequencing of kidneys from 7 weeks TRF plus FAN treatment reveals preservation of kidney function and metabolic homeostasis in injury. **(A)** Principal component analyses of vehicle and FAN treated kidneys. (n=4/group) **(B)** Volcano plot of differentially expressed genes (TRF-AL) using adjusted p.val < 0.05 and fold change > 2. **(C)** Violin plot of composite score of IRF8 gene cluster identified through Boolean analysis to predict sample identity in folic-treated samples. Top 10 pathways downregulated **(D)** and **(F)** heatmap of the mRNA expression of genes related to extracellular matrix protein, TGFβ and WNT signaling pathways that were downregulated in TRF. **(E)** Top 10 pathways upregulated by TRF in FAN model of kidney injury. Circle sizes in C and E indicate the number of genes in pathway and colored by enrichment score. **(G)** Network visualization of top pathways up and down regulated by TRF in FAN model of kidney injury. Immune related network of genes that are up regulated in ad lib fed mice (relative to TRF) are shaded with yellow background. Network of genes downregulated in TRF belonging to xenobiotics metabolism, general metabolism and mitochondria metabolism are shaded in red, green and blue respectively.

A total of 1,195 and 680 genes were up- and down-regulated respectively in FAN AL relative to FAN TRF group (Figure 4B), compared to 26 up- and 46 down-regulated genes observed in vehicle-treated mice under TRF compared to AL. However, the comparison of FAN AL vs Veh AL with FAN TRF vs FAN AL revealed that many genes induced by folic acid were modulated by the effect of TRF. More than half of the genes (54%, Supplementary Table 2) that were up-regulated following FAN treatment in the FAN-AL group relative to Veh AL were down-regulated in the FAN TRF group relative to FAN AL group. This suggests that TRF exerted a molecular brake to the FAN effect.

When this dataset was applied to the Boolean implication network constructed with human data, the network was able to accurately predict sample identity against AL and TRF (p=0.0208, Figure 4C). Additionally, genes in the previously identified IRF8 cluster were recapitulated in our FAN-injury model with 46% (32/69 genes) of these being significantly reduced by TRF (Figure S3A). TRF appeared to rescue expressions of *Lipa, Aoah, Sod2, Bid* and attenuated injury-induced genes like *Cd163, Tlr2*, and *Itgb2*.

Pathway enrichment analysis between FAN TRF vs FAN AL was performed using over-representation (ORA) and gene-set enrichment (GSEA) on DE genes, with a cutoff of adjusted p-value < 0.05 and a fold change > 2. The top pathways enriched in down-regulated genes in TRF were related to extracellular matrix organization, collagen metabolism, lymphocyte proliferation, and response to wounding (Figure 4D, E). Strikingly, the pathways enriched in the upregulated genes were involved almost exclusively in metabolic processes with the top pathways related to lipid metabolism, fatty acid metabolism, ATP synthesis, oxidative phosphorylation, and drug metabolism (Figure 4F). GSEA results were then visualized using Cytoscape plug-in, EnrichmentMap, revealing highly interconnected pathways. Specifically, immune activation (yellow) was distinctly upregulated in AL (Figure 4G). Three novel pathways emerging from the network enrichment suggested TRF modulated kidney metabolism (green), mitochondrial metabolism (blue), and retained drug metabolism (red) under FAN injury model (Figure 4G). To gain more insight, we also compared our data to MitoCarta3.0 (701/1140 genes FDR<0.05), an inventory of mammalian mitochondrial proteins, we find that TRF substantially increases expression of nuclear encoded mitochondrial genes and potentially modulates DNA repair and apoptosis. Furthermore, deconvolution of this bulk-RNA sequencing using available single cell datasets (GSE107585, GSE197266) found preservation of functional cells (proximal tubule/ distal convoluted tubule cells) and limited fibrotic (endothelial) and inflammatory (macrophages) cell proportions within the kidney in TRF groups compared to AL (Figure S3B).

Next, we used a targeted approach to assess how TRF affected the expression of key genes encoding critical steps in macronutrient metabolism. We found TRF increased the expression of genes implicated in fatty acid transport (*Cpt1a, Cpt2*) into the mitochondria for beta-oxidation (*Acadm, Hadh*) and reduced the expression of genes implicated in lipid synthesis (*Scd1, Fasn*). TRF also increased the expression of genes implicated in ketone metabolism (*Acat, Hmgcs2, Hmgcl*). Furthermore, TRF increased the expression of oxaloacetate (*Mdh1*) and pyruvate (*Pcx, Mdh2*) synthesis and changed the balance between gluconeogenesis and glycolysis by increasing gluconeogenesis (*Pck1, G6pc, Fbp1*), and decreasing glycolysis (*Hk1, Pfkl*) (Figure 5).

**Figure 5.**
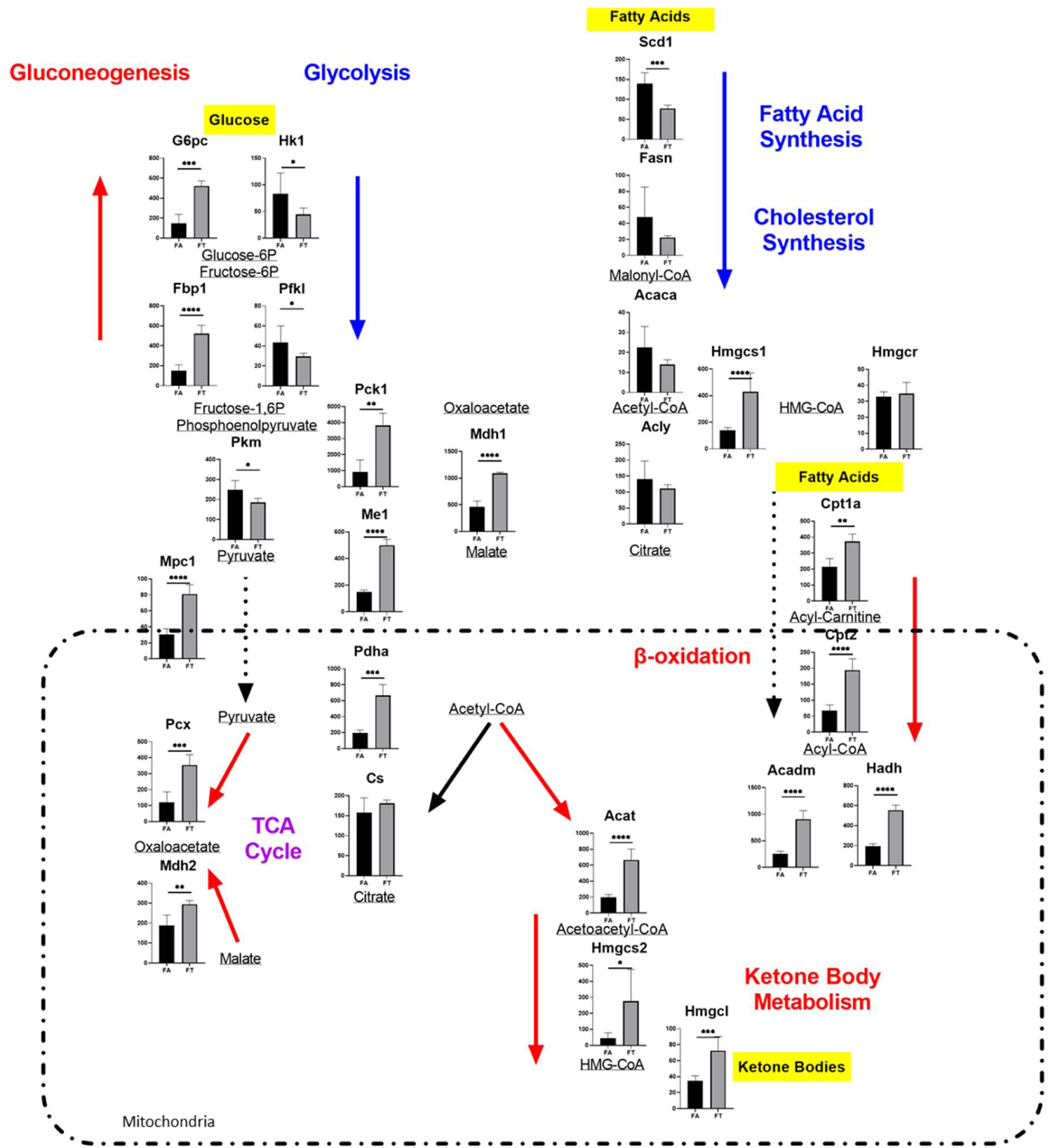
Targeted analysis demonstrates time-restricted feeding attenuates metabolic dysfunction associated with kidney damage and fibrosis. Gene expression of rate limiting/ irreversible reactions within metabolism. (n=4/group) The data represent the mean ± SEM. *P<0.05, **P<0.01, ***P<0.001 compared with Ad Libitum.

We used qPCR analysis to confirm these results in the UUO and short-term TRF FAN models as well. In agreement with the sequencing data, we found that mRNA expression of metabolism and mitochondrial-related genes correlated with the expression in mice, both under short-term TRF subjected to FAN or UUO (Figure S4A-C). Although, acetyl-CoA entry into the tricarboxylic acid (TCA) cycle (*Cs*) was not different between AL and TRF, the increased expression of pyruvate transporter (*Mpc1)* depicts increased conversion of pyruvate to oxaloacetate (*Pcx*) (Figure 5). Mice under the 1-week TRF protocol also showed increased expression of genes involved in fatty acid oxidation (*Cpt1a and Pparα*) and in gluconeogenesis (*Pcx, G6pc and Pepck*). Of note, the expression of *Pparα* and gluconeogenesis-related genes was also increased in control mice, reiterating the potential of TRF to enhance basal kidney function (Figure S4A). Conversely, mRNA expression of *Chrebp* and *Srebp1*, two transcription factors involved in lipogenesis (Linden et al., 2018; Ortega-Prieto & Postic, 2019) was significantly lower in mice on TRF subjected to kidney damage, and decreases were seen in the expression of the gene coding for fatty acid synthase (*Fasn*) in both models after 7 weeks (Figure S4B-C).

The protein expression of the mitochondrial transcription factor, TFAM, which plays a key role in the metabolic reprogramming associated with kidney fibrosis (Huang et al., 2018), was preserved in kidneys from mice on TRF for 1 or 7 weeks compared to AL mice, as demonstrated by immunohistochemistry (Figure S5A, B). Assessment of mitochondrial numbers through quantification of mitoDNA over nuclear DNA revealed an increase in mitochondrial copy number in kidneys from mice subjected to the 7 weeks TRF protocol (Figure S5C). Expression of mitochondrial-function related genes (*Tfam, Tomm20 and mtCytb*), were increased in all groups under TRF, particularly in the 7-weeks + FAN treatment (Figure S5D-F). Overall, these results suggest that TRF facilitated optimized FAO and protected mitochondrial health upon injury, contributing to the prevention of kidney fibrosis. Together, these data suggest an early protective role of TRF from metabolic derangement associated with kidney injury.

## Discussion

Obesity is a major risk factor for the development of diabetes, hypertension, and chronic kidney disease (Kopple, 2010; Whaley-Connell & Sowers, 2017). Numerous factors contribute to its progression, with aberrant feeding patterns emerging as a key risk. High-fat, high-sucrose diet has been widely used as a mouse experimental model for the induction of obesity, hypertension, and glucose intolerance. Since these pathologies are the main causes for CKD progression and have been previously shown to be ameliorated by TRF, we set out to determine whether TRF could prevent injury-induced damage. We utilized machine learning to test whether TRF could mitigate kidney injury. Through Boolean analysis, we anticipated TRF would alleviate inflammation-induced damage and fibrosis. Indeed, we observed a decrease in the expression of pro-inflammatory markers in TRF mice subjected to two different models of kidney injury. Moreover, we found that both short- and long-term periods of TRF were protective against injury-induced structural changes and fibrosis in the kidney, resulting in improved kidney function. TRF constituted an effective intervention that improves the robustness of circadian rhythms in metabolic tissues and results in beneficial metabolic outcomes (Chaix et al., 2019; Chaix et al., 2014; Duncan et al., 2016; Hatori et al., 2012; Sherman et al., 2012; Sundaram & Yan, 2016; Woodie et al., 2018).

After 6 weeks of high-fat feeding, AL mice weighed significantly more than TRF mice, with about 10% more body fat, typically manifest glucose intolerance, and demonstrated more severe kidney damage. Of note, even the short 1-week TRF protocol was still able to mitigate the damage induced by FAN, a model that is independent of weight difference or glucose intolerance, supporting the idea of an early protective effect of TRF that may be independent of acting through its effects on glucose intolerance. To our knowledge, this is among the first studies documenting the relevance of this chrono-nutrition intervention in CKD (Fang et al., 2023; Lao et al., 2023). Most recently, others have showed that TRF slowed the cyst progression in a rat model of polycystic kidney disease (PKD) (Torres et al., 2019), enhanced cell cycle regulation by reinforcing clock-dependent oscillations (Fang et al., 2023), and improved renal function in overweight and obese patients with stage 3-4 CKD (Lao et al., 2023). Furthermore, our meta-analyses of data from human CKD patients found genes associated with CKD were recapitulated in the FAN model and TRF was able to significantly attenuate the changes in expression of more than 40% of these genes towards levels found in healthy kidneys. Based on transcriptomic data, the two main proposed pathways by which TRF protects from inflammation/ fibrosis and preserves renal function are: 1) regulation of metabolic state and 2) mitigation of ER-stress and mitochondrial exhaustion.

The kidney is a highly metabolically active organ, rich in mitochondria to fulfill homeostatic conservation of the internal milieu through selective solute reabsorption and secretion of the glomerular filtrate. Renal tubule cells require high levels of ATP production, mostly supplied by oxidative metabolism. Abnormal mitochondrial or defective mitochondrial fatty acid oxidation in tubular epithelial cells is linked to ATP-depletion, lipotoxicity (Ge et al., 2020), apoptosis, fibrosis, and kidney dysfunction (Kang et al., 2015). Reciprocally, elevated FAO through renal tubule-specific overexpression of *Cpt1a* was demonstrated to protect from fibrosis (Miguel et al., 2021). Our study shows that kidneys from mice on TRF have higher expression of genes involved in fatty acid oxidation, ATP synthesis, and ketone metabolism together with increased mitochondrial number, and preemptively protect against insults. This shift towards alternative fuel utilization prompted during fasting periods likely also protect against lipotoxicity in the kidney from high-fat feeding. These observations are consistent with what we previously observed in the liver (Chaix et al., 2021; Chaix et al., 2014; Hatori et al., 2012).

Our results also showed an increased expression in gluconeogenesis-related genes in kidneys from mice under TRF, whereas opposite results were previously reported in livers (Hatori et al., 2012). The contribution to the rate of daily glucose synthesis by each organ depends on the nutritional balance: hepatic glucose production is the main endogenous glucose source during the post-absorptive state, whereas the kidney increases its glucose synthesis after long fasting periods (Kaneko et al., 2018; Mithieux et al., 2006; Owen et al., 1969). Proximal tubule epithelial cells are crucial for glucose reabsorption and for the maintenance of body-glucose levels(Eaton & Pooler, 2018). Importantly, kidney gluconeogenesis is exclusively performed in these cells, as they are the only ones that express the required enzymes (Mather & Pollock, 2011). Deconvolution of our bulk-sequencing estimates TRF preserves both proximal tubule and distal convoluted tubule epithelial cells while having reduced macrophage signatures. The transcriptional enrichment of gluconeogenic-related genes is evidence for an enhanced contribution of the kidney to gluconeogenesis, to maintain an adequate homeostasis of body-glucose under TRF conditions. This increase towards aerobic respiration is likely supported through utilization of available fatty acids, while generating ATP supply for filtration transport, and consequently prevents deposition of intracellular lipids that may interfere with healthy kidney function.

TRF also mitigates the response to injury by enhancing underlying circadian rhythms through oscillations reinforced by feeding and fasting, causing a switch between metabolic environments either conducive to early wound healing response or cellular maintenance. Heightened unfolded protein response and associated ER-stress have been correlated with the promotion of renal fibrosis in ischemic and nephrotoxic kidney injury (Chiang et al., 2011; Kim et al., 2015; Yan et al., 2018). Sustained ER stress has been shown to inhibit cell cycle progression and induce apoptosis upon irreversible damage (Hetz et al., 2020) ultimately leading to maladaptive repair and subsequently chronic inflammation and fibrosis (Yang et al., 2017). The amount of physiological damage prevented by TRF was not trivial as the effects were reflected in BUN and creatinine levels, which are clinical diagnostic markers of kidney function. We observed a decreased expression of ER-stress related genes in kidneys subjected to injury under 7 weeks of TRF. In addition, our histological observations showed a decrease in inflammation, less abnormal cell morphology and collagen deposition. These observations support a potential mechanism by which TRF prevents the establishment of inflammation and fibrosis by ameliorating UPR and preventing ER stress. Further work is required to test this hypothesis. Interestingly, the ability of TRF to maintain protein homeostasis has been documented in other tissues and models (Gill et al., 2015). Damage incurred by acute injury or chronic conditions typically triggers inflammation and disruption to the kidney molecular clock which can further exacerbate metabolic failure and development of fibrosis, frequently leading to organ dysfunction (Fang et al., 2023; Rey-Serra et al., 2023).

Circadian rhythm and metabolic pathways are reciprocally regulated, and hence metabolic processes can be coordinated with the feeding/fasting cycles (Stokkan et al., 2001). We recently showed that the molecular clock is a key proponent disrupted by renal damage and promotes further metabolic derangement leading to fibrosis and CKD (Rey-Serra et al., 2023). Here, we propose that TRF enhances oscillations of metabolic genes and regulation of cell cycle/ proliferation through the imposed feeding/ fasting cycle to prioritize FAO when supporting activity and repair/ proliferation mechanisms during inactivity. Thus, TRF represents a chrono-dietary modification, capable of shielding from inflammation in different pathological settings and can be considered potent intervention for human CKD patients. Overall, our results support that time-restricted feeding is a preventative lifestyle intervention for inflammation, fibrosis and metabolic dysfunction associated with/ attributed to kidney damage.

Limitations of the study: The study was done on young male mice only, while CKD is known to affect both sexes and at an older age. The disease models used in this study accelerate disease onset and progression. The Boolean implication network analyses identified five clusters of genes associated with CKD, of which only one was affected by TRF. While this approach validated the predictive value of the model, it also leaves open the possibility that additional interventions alone or in combination with TRF can be more effective in preventing or managing CKD.

## Acknowledgements

This publication includes data generated at the UC San Diego IGM Genomics Center utilizing an Illumina NovaSeq 6000 that was purchased with funding from a National Institutes of Health SIG grant (#S10 OD026929). This work used Jetstream2 at Indiana University through allocation BIO220036/ BIO210090 from the Extreme Science and Engineering Discovery Environment (XSEDE), which was supported by National Science Foundation grant number #1548562. Research in the laboratory of SP is supported by NIH grants CA258221, and AG068550, the Wu Tsai Human Performance Alliance and the Joe and Clara Tsai Foundation. Experiments and analyses utilized the Salk Institute core services supported by NIH grant P30CA014195. This work was supported by a grant from the Ministerio de Ciencia e Innovación PID2019-104233RB-100/AEI/10.13039/501100011033 (SL). Carlos Rey-Serra has been the recipient of an FPI research-training contract from the Spanish Research State Agency (BES-2016-076735). AC is supported by AHA grant 18CDA34110292 and NIA grant 5R01AG065993.

---

**Supplementary Figure S1.**
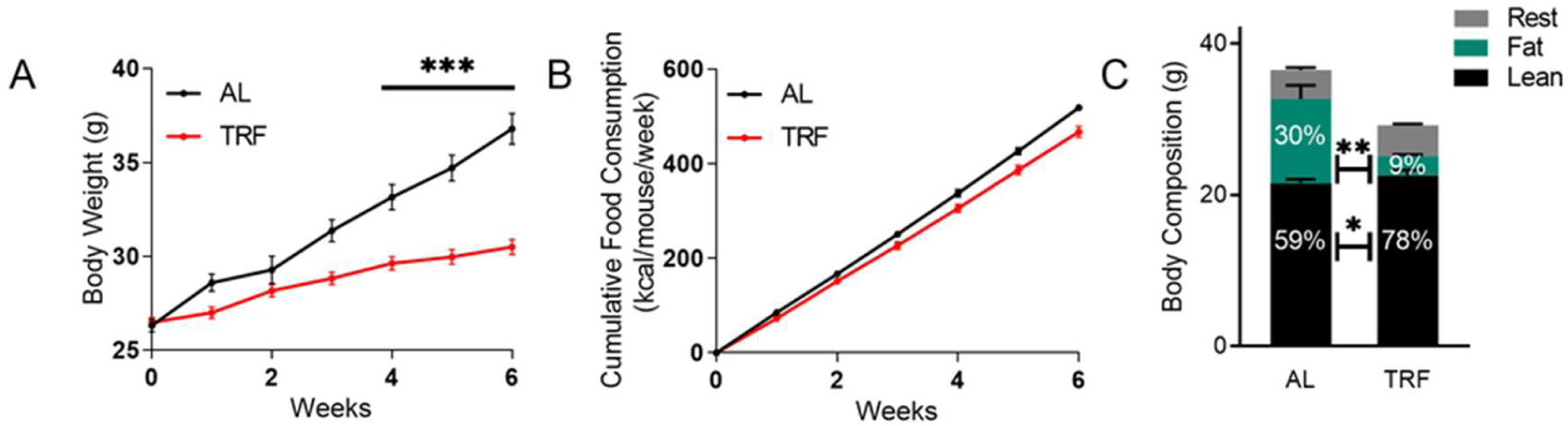
Body weight, food consumption, and body composition. **(A)** Body weight of male C57BL6 mice fed 45% HFD over 6 weeks under Ad Libitum (AL) or time-restricted feeding. (n=30/group) **(B)** Food consumption measured as kcal/mouse/week. **(C)** Body composition in grams with calculated percentage in relation to total body weight. (n=8/group).

**Supplementary Figure S2.**
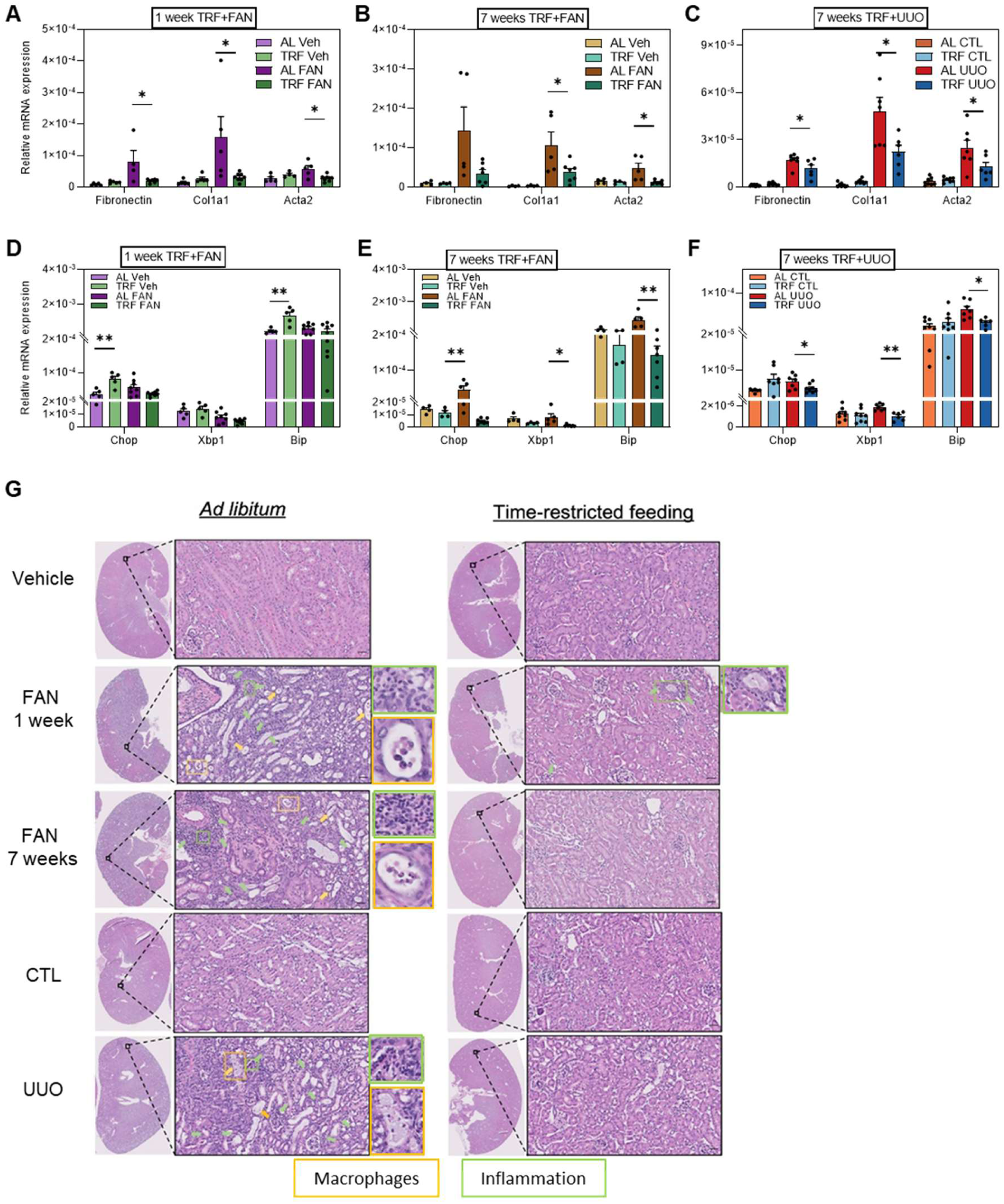
Kidney inflammation in FAN- and UUO-induced kidney damage is reduced upon time-restricted feeding. mRNA expression of fibrotic and mitochondrial genes in **(A, D)** 1-week TRF plus FAN, **(B, E)** 7-weeks TRF plus FAN, and **(C, F)** 7-weeks TRF plus UUO. **(G)** Representative microphotographs of H&E staining of vehicle-treated and folic acid-treated kidneys from 1-week and 7-weeks TRF plus FAN, and contralateral/ obstructed kidneys from 7-weeks TRF plus UUO protocol. Arrows/rectangles denote interstitial inflammation (green), macrophages in the tubular lumen (yellow). Scale bar: 50 µm.

**Supplementary Figure S3.**
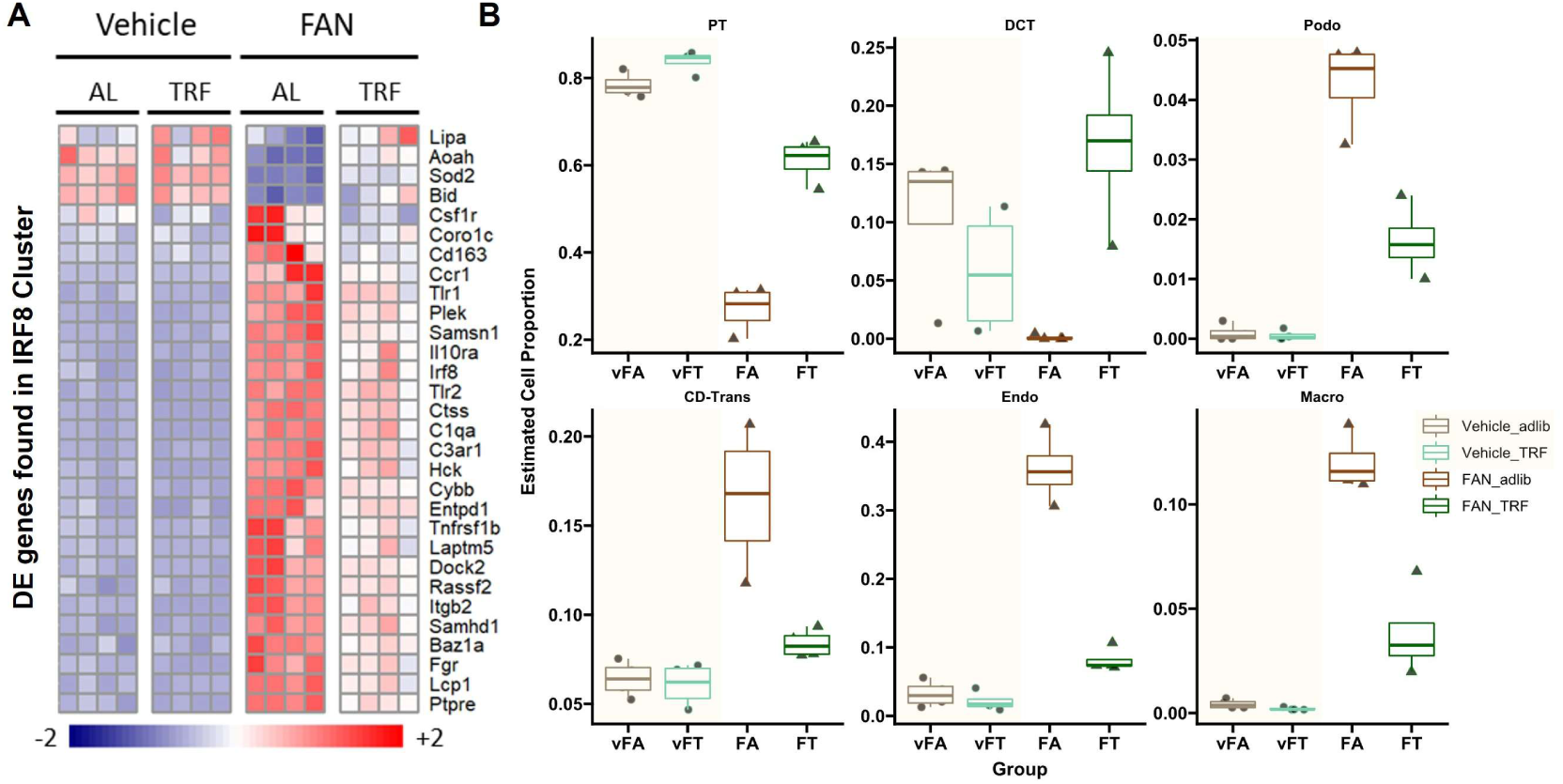
Deconvolution of bulk RNA-sequencing shows TRF attenuates injury-induced inflammation and maintains kidney cell population. **(A)** Heatmap of Boolean implicated genes in human CKD that were significantly altered by TRF in 7 weeks plus FAN treatment (n=4/group). **(B)** Estimated cell proportion in seven-weeks TRF plus two-weeks FAN bulk-RNA sequencing deconvoluted from single cell dataset GSE107585. Proportion of estimated cell types: PT: proximal tubule, DCT: distal convoluted tubule, Podo: podocyte, CD-Trans: collecting duct-transitional cell, Endo: endothelial, Macro: macrophages.

**Supplementary Figure S4.**
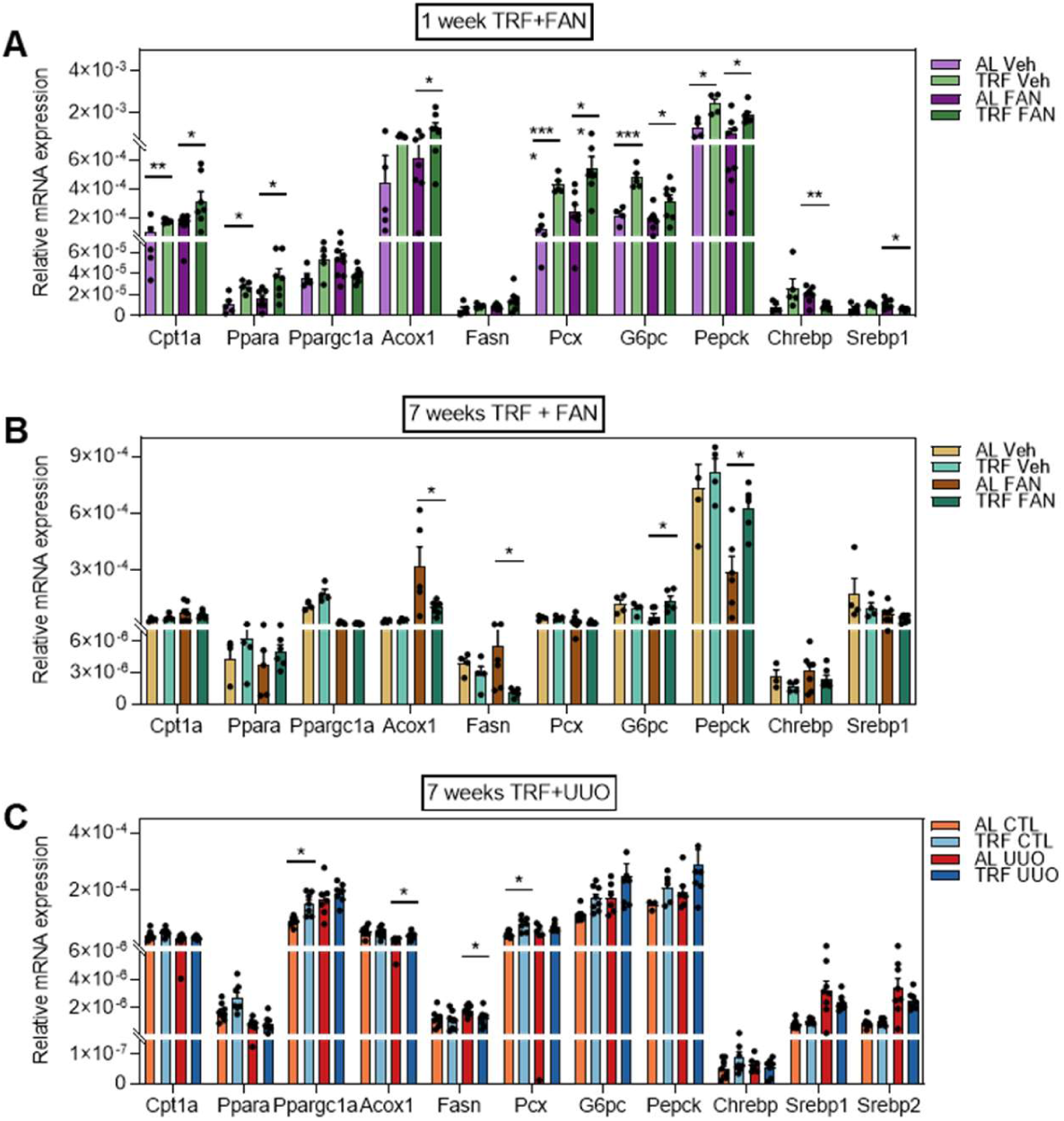
qPCR validation of metabolic gene signature and bulk RNA deconvolution. mRNA expression of lipid-metabolic genes in mice under **(A)** one-week TRF plus two-weeks FAN, **(B)** seven-weeks TRF plus two-weeks FAN, and **(C)** seven-weeks TRF plus three-days UUO. (n=6-8/group).

**Supplementary Figure S5.**
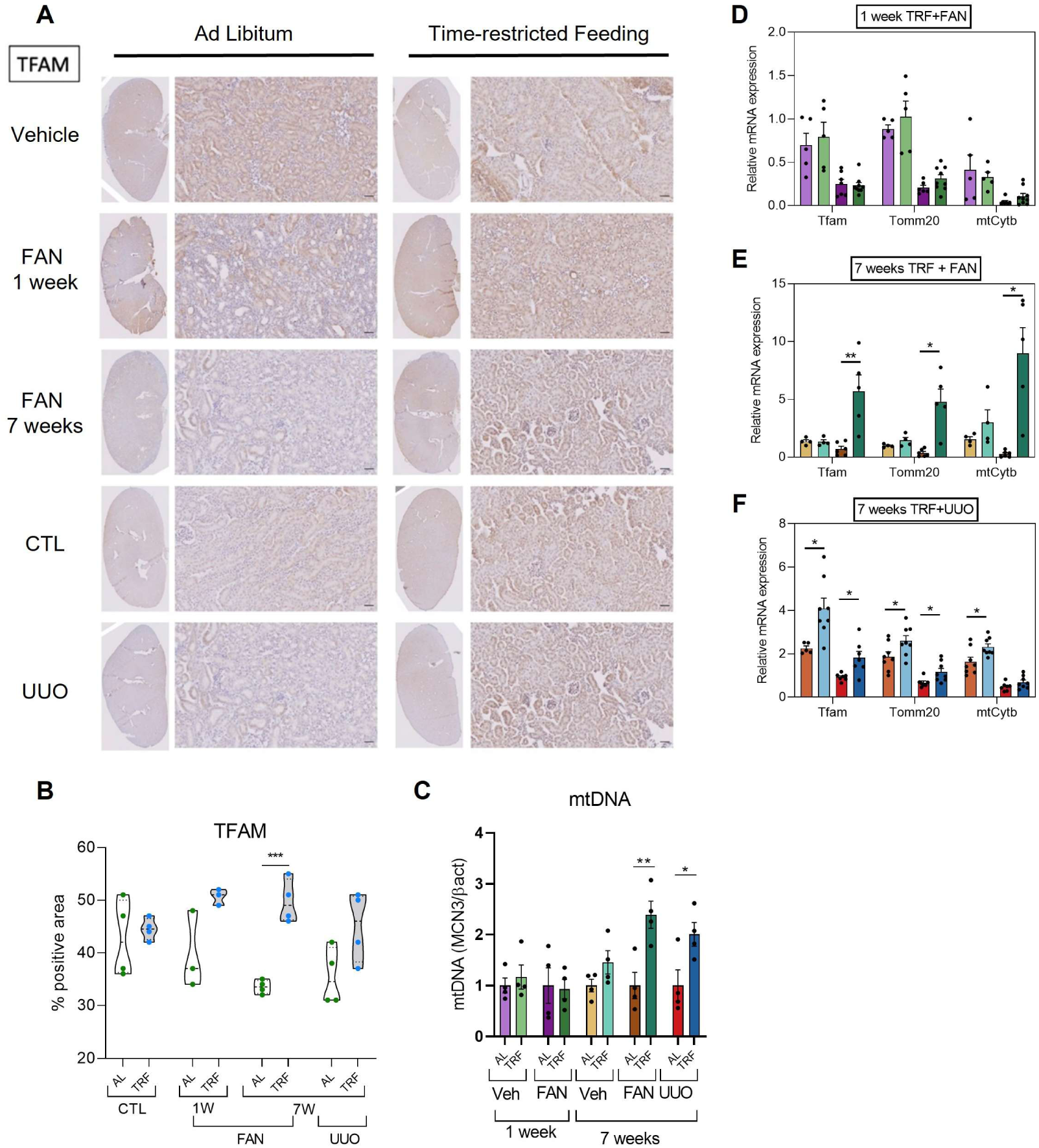
Mitochondrial function is preserved under TRF during kidney injury. **(A)** Representative pictures of TFAM IHC staining of kidneys from mice that had been under one or seven weeks of TRF or AL regimens in the 2-weeks FAN (vehicle-treated and folic acid-treated mice) or 3 days-UUO models. Scale bar: 50 *µm.* **(B)** Quantification of TFAM IHC in vehicle and folic acid-treated kidney sections FAN and UUO models. (n=4/group) **(C)** Mitochondrial DNA copy number quantification in kidney (n=4/group). mRNA expression of mitochondrial genes in mice under **(D)** one-week TRF plus two-weeks FAN, **(E)** seven-weeks TRF plus two-weeks FAN, **(F)** seven-weeks TRF plus three-days UUO.

**Supplementary Table S1.** Genes with conserved expression in Boolean cluster IRF8. Of the 69 genes in the human IRF8 Boolean cluster, 60 were reliably detected in mouse kidney transcriptomes across all samples in this study. This table lists the names of those 60 genes along with summary statistics of their expression differences between ad libitum–fed and time-restricted–fed (TRF) mice treated with folic acid.

**Supplementary Table S2.** Normalized gene expression values and summary statistics of kidney transcriptome from Folic acid or vehicle treated mice. Metadata and TMM normalized gene expression values are shown. The results of dSeq2 differential gene expression analyses and the respective statistics including false discovery rates for comparison between folic acid adlib and folic acid TRF mice are presented.

**Supplementary Table S3. Quantitative-PCR primers of various mouse genes assessed in this study.** The primer sequences are designed against reference mouse genome.

