## Supplementary figures for "Time-Restricted Feeding Attenuates Kidney Damage and Preserves Renal Function in Mouse Model of Chronic Kidney Disease"

### Supplementary Figure S1

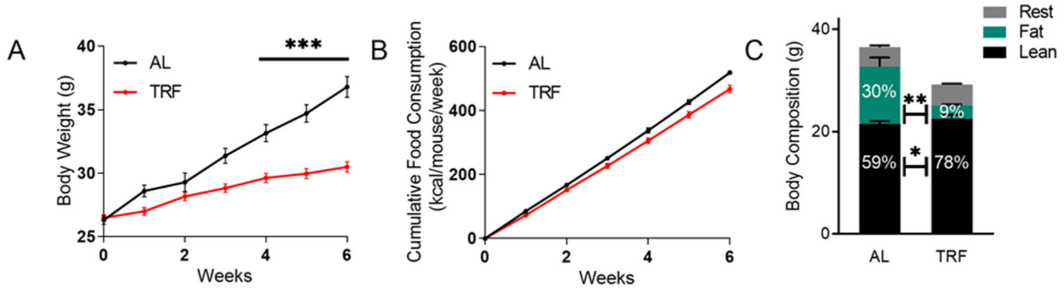


**Supplementary Figure S1. Body weight, food consumption, and body composition. (A)** Body weight of male C57BL6 mice fed 45% HFD over 6 weeks under Ad Libitum (AL) or time- restricted feeding. (n=30/group) **(B)** Food consumption measured as kcal/mouse/week. **(C)** Body composition in grams with calculated percentage in relation to total body weight. (n=8/group).

### Supplementary Figure S2

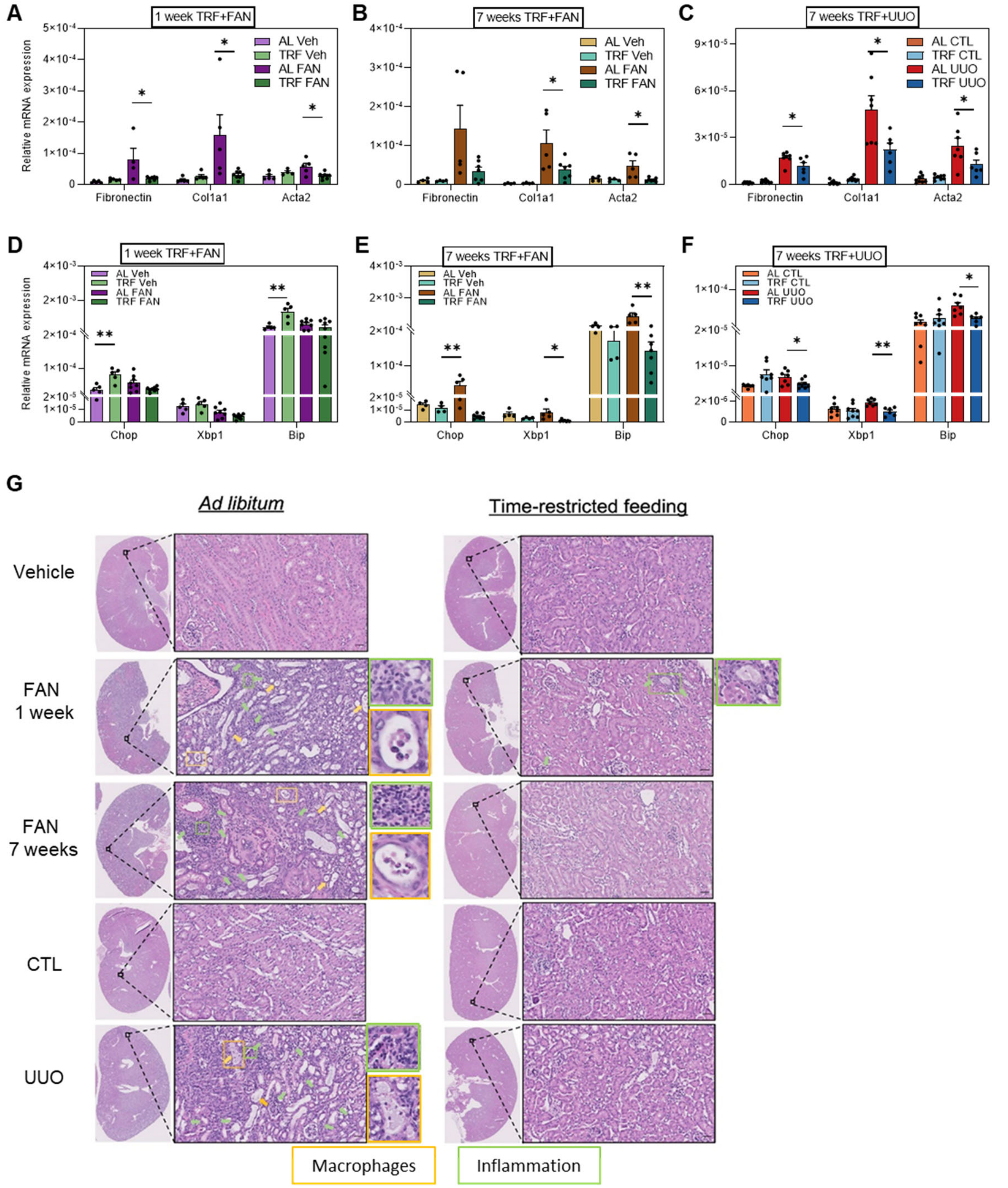


**Supplementary Figure S2. Kidney inflammation in FAN- and UUO-induced kidney damage is reduced upon time-restricted feeding.** mRNA expression of fibrotic and mitochondrial genes in **(A, D)** 1-week TRF plus FAN, **(B, E)** 7-weeks TRF plus FAN, and **(C, F)** 7-weeks TRF plus UUO. **(G)** Representative microphotographs of H&E staining of vehicle-treated and folic acid-treated kidneys from 1-week and 7-weeks TRF plus FAN, and contralateral/ obstructed kidneys from 7-weeks TRF plus UUO protocol. Arrows/rectangles denote interstitial inflammation (green), macrophages in the tubular lumen (yellow). Scale bar: 50 μm.

#### Supplementary Figure S3

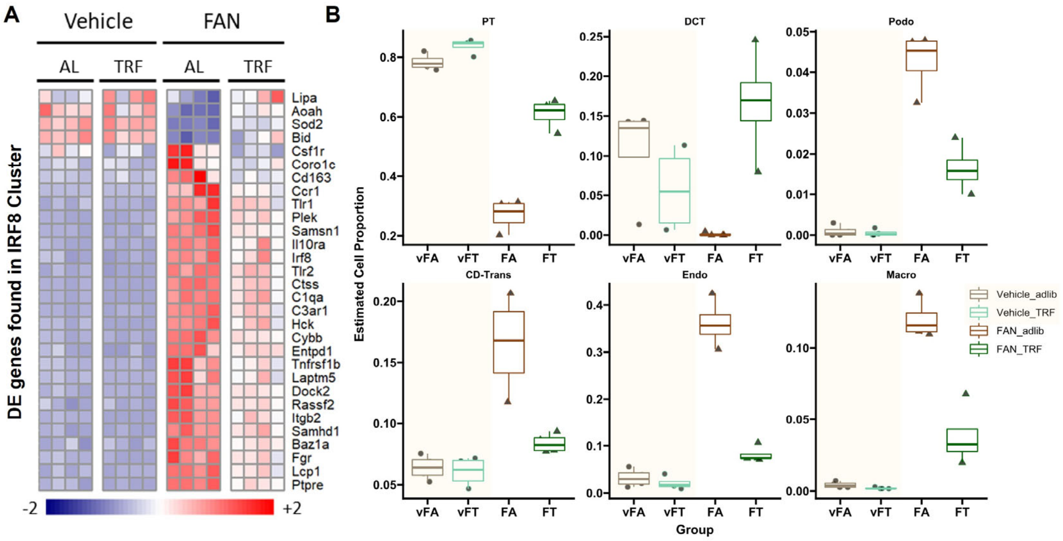


**Supplementary Figure S3. Deconvolution of bulk RNA-sequencing shows TRF attenuates injury-induced inflammation and maintains kidney cell population. (A)** Heatmap of Boolean implicated genes in human CKD that were significantly altered by TRF in 7 weeks plus FAN treatment (n=4/group). **(B)** Estimated cell proportion in seven-weeks TRF plus two-weeks FAN bulk-RNA sequencing deconvoluted from single cell dataset GSE107585. Proportion of estimated cell types: PT: proximal tubule, DCT: distal convoluted tubule, Podo: podocyte, CD-Trans: collecting duct- transitional cell, Endo: endothelial, Macro: macrophages.

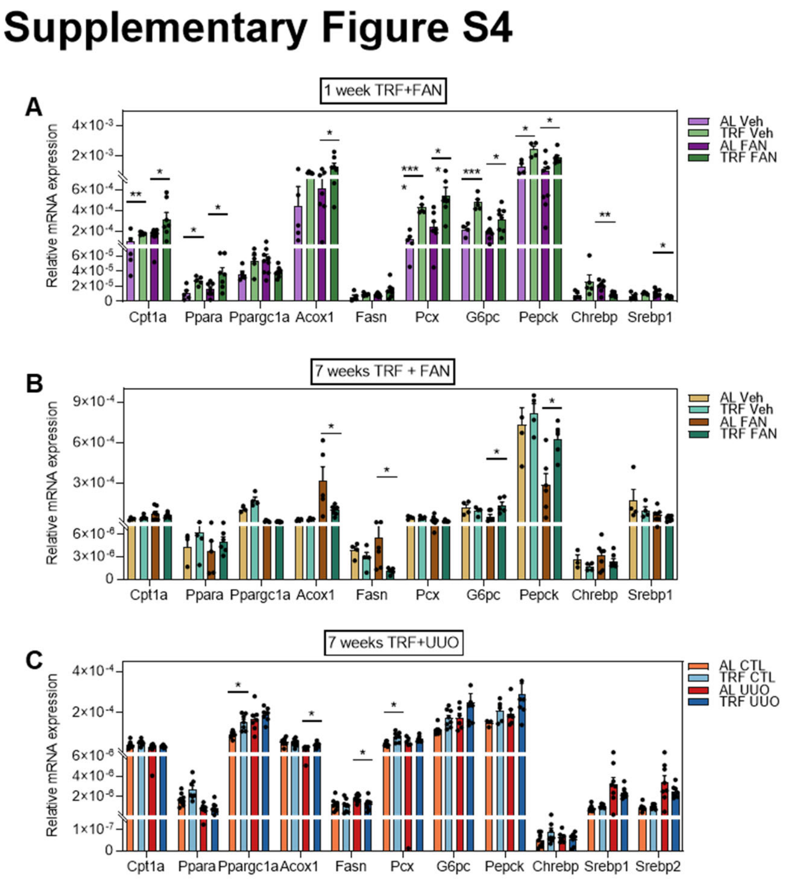


**Supplementary Figure S4. qPCR validation of metabolic gene signature and bulk RNA deconvolution.** mRNA expression of lipid-metabolic genes in mice under **(A)** one-week TRF plus two-weeks FAN, **(B)** seven-weeks TRF plus two-weeks FAN, and **(C)** seven-weeks TRF plus three-days UUO. (n=6-8/group).

### Supplementary Figure S5

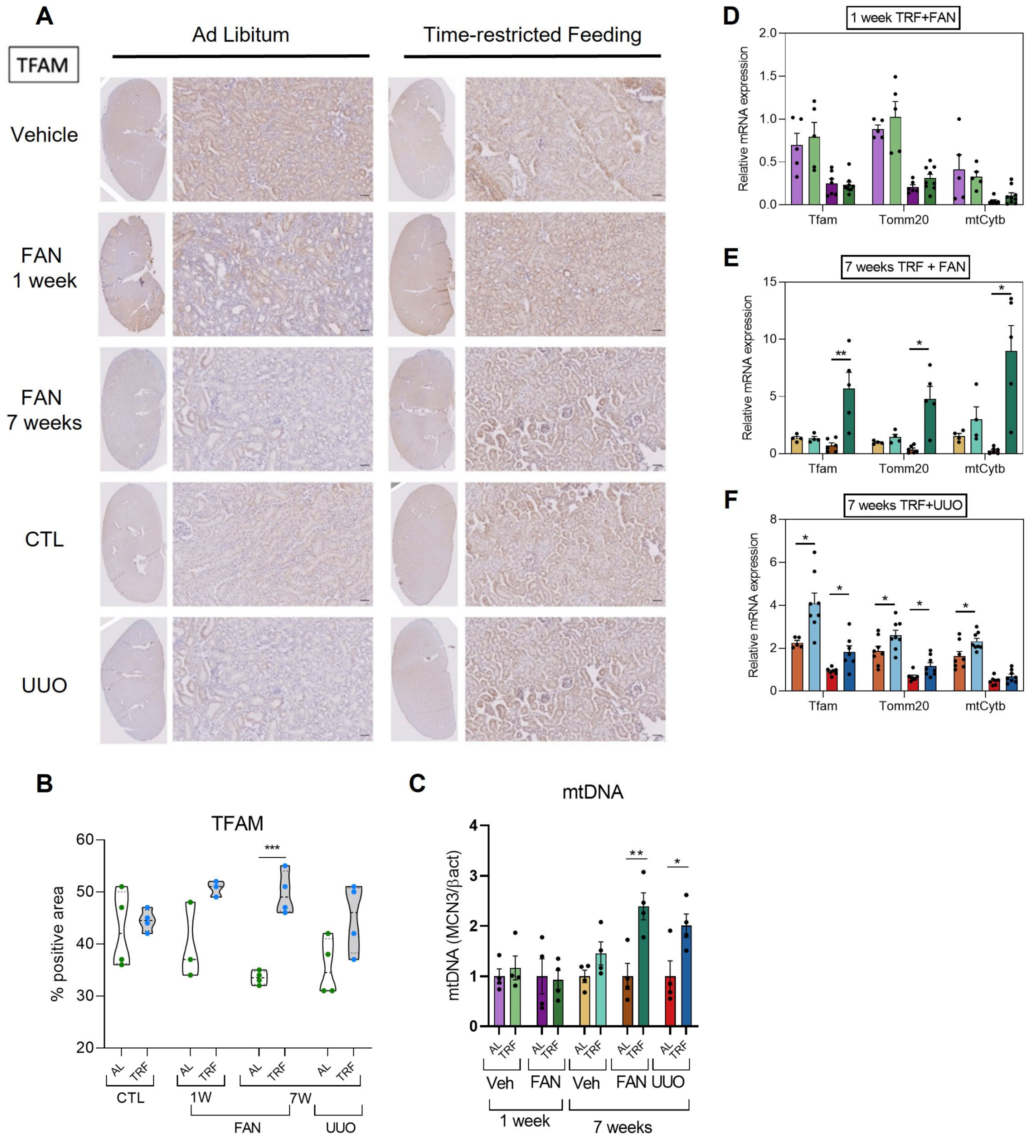


**Supplementary Figure S5. Mitochondrial function is preserved under TRF during kidney injury. (A)** Representative pictures of TFAM IHC staining of kidneys from mice that had been under one or seven weeks of TRF or AL regimens in the 2-weeks FAN (vehicle-treated and folic acid-treated mice) or 3 days-UUO models. Scale bar: 50 *μm.* **(B)** Quantification of TFAM IHC in vehicle and folic acid-treated kidney sections FAN and UUO models. (n=4/group) **(C)** Mitochondrial DNA copy number quantification in kidney (n=4/group). mRNA expression of mitochondrial genes in mice under **(D)** one-week TRF plus two-weeks FAN, **(E)** seven-weeks TRF plus two-weeks FAN, **(F)** seven-weeks TRF plus three-days UUO.

**Supplementary Table S1.** Genes with conserved expression in Boolean cluster IRF8. Of the 69 genes in the human IRF8 Boolean cluster, 60 were reliably detected in mouse kidney transcriptomes across all samples in this study. This table lists the names of those 60 genes along with summary statistics of their expression differences between ad libitum–fed and time-restricted–fed (TRF) mice treated with folic acid.

**Supplementary Table S2.** Normalized gene expression values and summary statistics of kidney transcriptome from Folic acid or vehicle treated mice. Metadata and TMM normalized gene expression values are shown. The results of dSeq2 differential gene expression analyses and the respective statistics including false discovery rates for comparison between folic acid adlib and folic acid TRF mice are presented.

**Supplementary Table S3. Quantitative-PCR primers of various mouse genes assessed in this study.** The primer sequences are designed against reference mouse genome.
